# Modulator Resolves Transcript-, Allele-, and Condition-associated RNA Modification Differences from Direct RNA Sequencing

**DOI:** 10.64898/2026.10.15.759385

**Authors:** Theodore M. Nelson, Michael Goneos, Caroline J. Aufgebauer, Paul Collier, Jacqueline Proszynski, Krista Ryon, Evan M. Violette, Eyup Alper Saygili, Nadia Houerbi, Stacy M. Horner, Christopher E. Mason

**Affiliations:** Department of Systems and Computational Biomedicine, Weill Cornell Medicine, New York, NY 10065, USA The HRH Prince Alwaleed Bin Talal Abdulaziz Alsaud Institute for Computational Biomedicine, Weill Cornell Medicine, New York, NY 10065, USA; Department of Molecular Genetics and Microbiology, Duke University School of Medicine, Durham, NC 27710, USA; Department of Integrated Immunobiology, Duke University School of Medicine, Durham, NC 27710, USA; Department of Medicine, Duke University School of Medicine, Durham, NC 27710, USA; The WorldQuant Initiative for Quantitative Prediction, Weill Cornell Medicine, New York, NY, 10065, USA; School of Medicine, Acibadem University, Istanbul, 34752, Turkey; The Feil Family Brain and Mind Research Institute, Weill Cornell Medicine, New York, NY, 10065, USA

## Abstract

Nanopore direct-RNA sequencing (DRS) reports RNA modifications on single molecules, but there is no comprehensive algorithm for processing, aligning, and differentiating base modifications across reads that are often 5′-truncated. Here we present Modulator, which assigns every read, truncated or not, to a “fragmentform,” defined by a shared transcript 3′ end and the splice-junctions upstream of it. Of note, Modulator builds fragmentforms from genomic alignments without a reference transcriptome, estimates per-base modification stoichiometry within each fragmentform, and classifies each stoichiometry difference as transcript-, allele-, or condition-dependent. In the control HG002 transcriptome, Modulator built 23,746 fragmentforms, with 15,620 candidate novel isoforms that carried poly(A) signals as often as previously annotated transcripts. Also, assigning 5′-truncated reads to fragmentforms nearly doubled the number of testable and differentially modified sites. m6A sites within 100 nucleotides (nt) of a splice junction or within 1 kb of a poly(A) site were less modified in fragmentforms that use that junction or poly(A) site, indicating that local transcript context shapes where m6A is deposited. From the same reads, Modulator called SNPs with 93% precision and no phasing errors against the Genome in a Bottle benchmark. It identified 131 SNPs that create or remove a modifiable nucleotide and 1,358 allele-specific modification differences, most involving SNPs more than 50 nt from the modified site. In cells infected with Zika virus, which remodels gene expression relative to uninfected cells, Modulator identified 1,253 transcript-independent modification stoichiometry changes. Of the m6A changes at sites measured by GLORI, 96% agreed in direction with single-nucleotide-resolution GLORI maps of the same samples. Thus, Modulator accurately separates changes in RNA modification from transcript-associated and allele-associated differences and can improve the utility of nanopore DRS.

**Availability and Implementation:** Modulator is free and open source: https://github.com/Theo-Nelson/Modulator. It is written in Python and runs on Linux OS.

## INTRODUCTION

An observed difference in RNA modification stoichiometry at a genomic position can reflect altered modification within a transcript, a shift in which transcripts carry the position, or allele-specific sequence variation. Current analyses cannot reliably distinguish among these possibilities. A genomic position may be represented in multiple transcripts from the same gene, and these transcripts can be modified at different stoichiometries (Gleeson *et al*. 2025). Alternative transcription start sites, alternative splicing, and 3′-end choice can each alter the transcript context surrounding a potentially modified nucleotide and influence how often it is modified (Tang *et al*. 2024). Genetic variation can also affect modification, either by creating or removing a modifiable nucleotide or by altering modification at a nearby site (Park and Cenik 2025). Consequently, a change measured across a genomic locus does not necessarily represent a change in modification within any individual transcript or allele.

Nanopore direct-RNA sequencing (DRS) offers a potential solution because it sequences native RNA molecules and preserves transcript-architecture and base modification information on individual reads (Jain *et al*. 2022, Esfahani *et al*. 2025). In principle, these measurements make it possible to determine the transcript and allele carrying each modified nucleotide, as well as their phase, and thus resolve the nature of a stoichiometry difference. Existing analyses, however, either measure modification at the genomic locus, thereby merging reads from different transcripts and alleles, or assign reads to transcript models inferred from a reference annotation or transcriptome assembly (Gleeson *et al*. 2025). The first approach obscures molecular heterogeneity, whereas the second depends on accurate transcript assignment. Although long-read transcriptome assembly and quantification tools can infer sample-specific transcript structures, most transcript-resolved modification analyses still require reads to be assigned to complete transcript models. Because DRS sequences RNA molecules from their 3′ ends toward their 5′ ends, many reads terminate before reaching the transcript 5′ end, preventing confident assignment to a full-length isoform and limiting the fraction of molecules that can be analyzed in transcript context (Depledge *et al*. 2019, Drexler, Choquet, and Churchman 2020, Mulroney *et al*. 2022). Many transcripts also remain unannotated even in well-studied organisms, and reference annotations may poorly represent transcripts expressed in particular cell types, conditions, disease states, or non-model organisms (Chen *et al*. 2023).

Although truncated DRS reads do not capture complete transcript structures, they retain two features that can define the local transcript context of a modified position: the transcript 3′ end, marked by its poly(A) tail, and the chain of splice junctions between the read’s 5′ endpoint and that 3′ end (Depledge *et al*. 2019, Dar *et al*. 2024). We reasoned that these shared features could be used to group reads without requiring knowledge of the transcript 5′ end. We define such a group as a fragmentform: the set of reads that share a transcript 3′ end and the same upstream splice-junction chain over the region represented by the reads. A read that extends only partway through the transcript can still be assigned to a fragmentform defined by its observed 3′ end and junction structure. Thus, a fragmentform does not claim to represent a complete transcript isoform. Instead, it captures the transcript architecture that is observable and relevant to a given read, allowing both truncated and full-length molecules to contribute to transcript-context-dependent modification analysis and stoichiometry.

Here we present Modulator, a reference-transcriptome-free framework that constructs fragmentforms from genomic alignments and assigns every read to the fragmentform its own features define. Within each fragmentform, Modulator estimates modification stoichiometry and tests whether that stoichiometry differs among fragmentforms from the same gene or locus. This approach distinguishes a modification difference associated with transcript architecture from one caused solely by a change in the abundance of transcripts carrying the site. By building fragmentforms directly from the reads rather than from a reference annotation, Modulator can also identify sample-specific combinations of novel splice junctions and 3′ ends while retaining reads that cannot be assigned to complete isoforms.

In addition, Modulator calls and phases single-nucleotide polymorphisms from the same DRS reads. Each RNA molecule can therefore be linked to its fragmentform, allele, and modification state. Long-read methods have established frameworks for connecting genotype with transcript structure, including allele-specific expression and splicing analyses (Fu *et al*. 2026, Huang *et al*. 2026), while DRS has revealed allele-specific m6A at the gene level (Park and Cenik 2025). These approaches link alleles to either transcript architecture or RNA modification, but do not jointly resolve transcript structure, genotype, and modification state on the same molecules. Integrating these measurements allows Modulator to identify allele-dependent modifications on the same molecules that define transcript context.

We apply Modulator to the HG002 transcriptome and to Zika virus infection, a condition that extensively remodels gene expression. These analyses show that including 5′-truncated reads substantially expands the set of modification sites that can be tested in a defined transcript context. They further reveal modification differences associated with splice-junction and poly(A)-site usage, genetic variation, and cellular condition. By separating modification changes from changes in the transcripts and alleles that carry the modified positions, Modulator provides a framework for determining the molecular source of RNA modification differences from DRS data and offers a rich annotation of nanopore DRS data.

## Results

### Fragmentforms recover transcript-context information from truncated reads

Direct RNA sequencing reads molecules from the 3′ end toward the 5′ end, and many reads terminate before reaching the 5′ end (Depledge *et al*. 2019, Drexler, Choquet, and Churchman 2020, Mulroney *et al*. 2022). We first assessed how often reads reached annotated transcript 5′ ends, assuming that the hg38 reference transcriptome accurately reports those ends. HG002 RNA was sequenced on two flow cells, designated library 1 and library 2. We restricted this analysis to the 7.7 million HG002 reads that matched annotated transcripts by junction chain and 3′ end. We defined a read as reaching the transcript 5′ end if its own 5′ end was within 50 nucleotides (nt) of the annotated end. For transcripts under 1 kb, 88% of reads reached the 5′ end, compared with 35% for transcripts over 5 kb (**Figure 1a**). Thus, as expected, longer transcripts exhibited more 5′ truncation than shorter transcripts.

**Figure 1.**
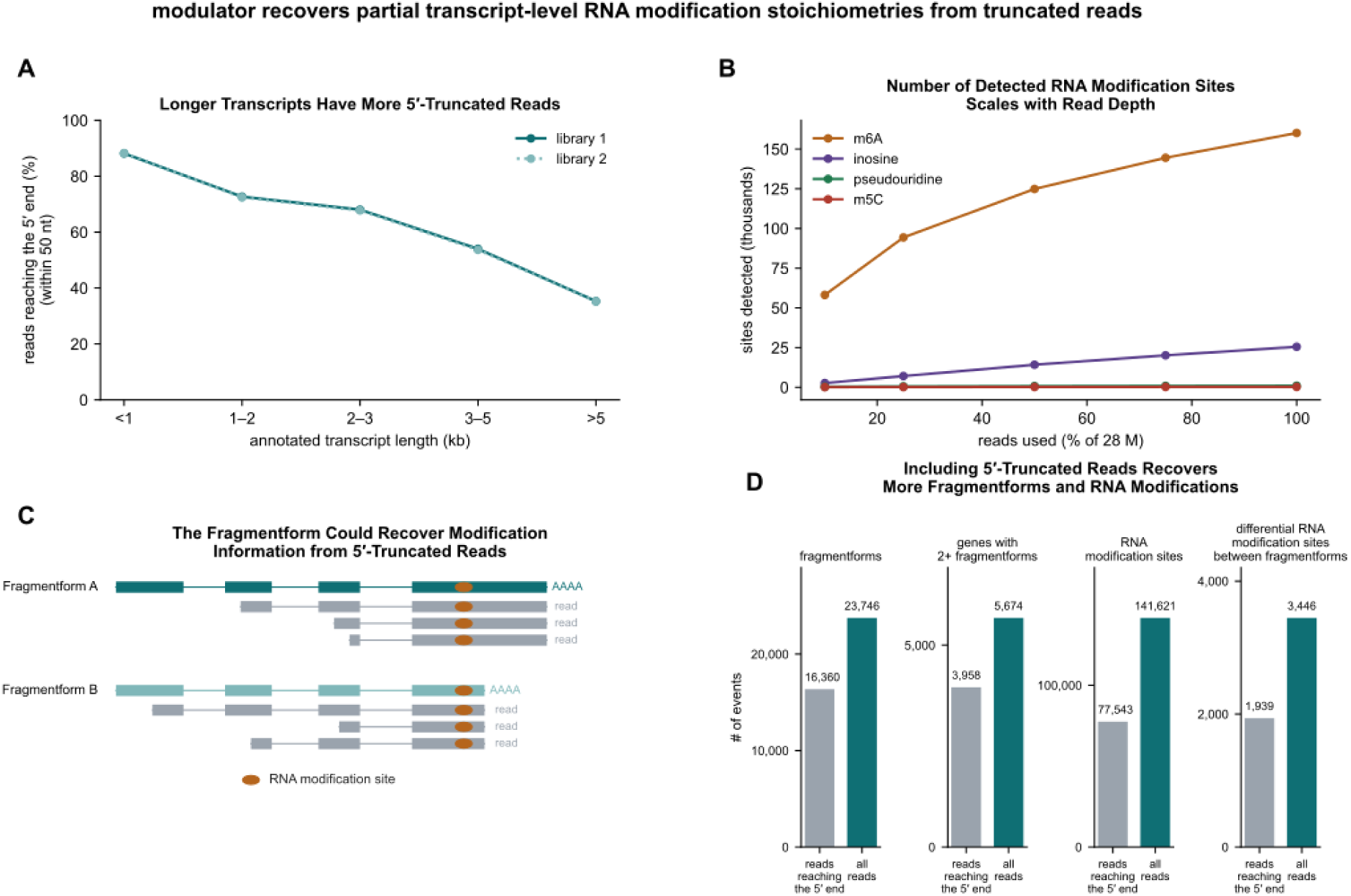
Modulator recovers partial transcript-level RNA modification stoichiometries from truncated reads. **(a)** *Longer transcripts have more 5′-truncated reads.* To measure how often a read reaches the 5′ end, we used the 7.7 million HG002 reads on fragmentforms that match a transcript within the hg38 reference transcriptome based on 3’ end and splice junction usage. The plot shows the percentage of reads whose own 5′ end is within 50 nt of the annotated 5′ end, based on the transcript length reported by the hg38 reference transcriptome. The HG002 RNA was sequenced twice on two different flow cells, labeled as ‘library 1’ and ‘library 2’. Of the reads on transcripts under 1 kb, 88% reach the 5′ end. Of the reads on transcripts over 5 kb, only 35% reach the 5’ end. **(b)** *The number of detected RNA modification sites scales with read depth.* For this experiment, the HG002 reads from the two different flow cells (‘library 1’ and ‘library 2’) were combined into a single sequencing dataset with 28.2 million reads mapped to the hg38 genome. We then took random subsets and measured the number of sites per modification. A site requires at least 20 reads and greater than 10% modified reads, Nfail + 1 ≤ Nmod, and Ndiff ≤ 3 × Nvalid_cov. m6A sites rise from 58,160 when 10% of available reads are used to 160,104 at 100% of available reads. **(c)** *The fragmentform could recover modification information from 5′-truncated reads*. Schematic. Two fragmentforms of one gene with three example reads shown in grey. Every read spans at least one splice junction, as Modulator requires by default. Each read is assigned to a fragmentform from its 3′ end and its junction chain, irrespective of 5’ truncation. The amber circle marks the demonstrative RNA modification site. **(d)** *Including 5′-truncated reads recovers more fragmentforms and RNA modifications*. Modulator was run with identical settings on two read sets: the 45% of HG002 reads whose 5′ end lies within 50 nt of an annotated 5′ end (grey) and all HG002 reads (dark teal). The four bar charts show the number of fragmentforms (left), genes with two or more fragmentforms (left, middle), RNA modification sites testable between fragmentforms (middle, right), and differential RNA modification sites between fragmentforms (right).

We next examined how sequencing depth affects the number of detectable RNA modification sites. modkit aggregates per-read calls at genomic positions and calculates modification stoichiometry at each site. We combined the two flow cells into a dataset containing 28.2 million reads mapped to hg38, then analyzed random subsets (10, 25, 50, 75, and 100%) of these reads. We counted sites with at least 20 reads and greater than 10% modification stoichiometry. The number of m6A sites rose from 58,160 at 10% of the reads to 160,104 at 100% of the reads, with no sign of a plateau (**Figure 1b**). Inosine sites rose from 2,674 to 25,547, pseU sites from 378 to 990, and m5C sites from 127 to 280 (**Figure 1b**). Site detection continued to increase with sequencing depth, motivating an approach that retains informative 5′-truncated reads.

To retain 5′-truncated reads, we developed the “fragmentform.” A fragmentform is defined by the 3′ end of a transcript and the splice junctions upstream of that end. We refer to the ordered list of these junctions as the “junction chain.” Because the nanopore reads each molecule from the 3′ end, every read contains the 3′ end of its molecule, and every spliced read contains a junction chain. Modulator assigns each read to a fragmentform from the 3′ end and junction chain of the read itself, so the read does not need to reach the 5′ end (**Figure 1c**). Modulator then compares the stoichiometry at each site between the fragmentforms of a gene. In summary, the fragmentform allows every read, 5′-truncated or not, to contribute to a transcript-level measurement of stoichiometry.

To measure the contribution of 5′-truncated reads directly, we ran Modulator twice on the same two HG002 libraries with identical settings: once using only reads reaching an annotated 5′ end (45% of the reads in each library) and once using all reads. With all reads, Modulator built 23,746 fragmentforms compared to 16,360 with only the reads reaching the 5′ end (**Figure 1d**). It found 5,674 genes with two or more fragmentforms compared to 3,958 (**Figure 1d**). It found 141,621 RNA modification sites testable between fragmentforms compared to 77,543, and 3,446 sites that differed between fragmentforms compared to 1,939 (**Figure 1d**). Thus, retaining 5′-truncated reads nearly doubled the number of testable and differential modification sites identified by Modulator.

### Evidence supporting fragmentform 3′ ends and abundance estimates

We next evaluated the evidence supporting the inferred fragmentforms. We first compared each fragmentform with the hg38 RefSeq annotation. Modulator classifies each fragmentform by its junction chain and its 3′ end (**Figure 2a**). An “exact match” has the same junction chain as an annotated transcript and a 3′ end within 25 nt of the annotated end. A “new junction chain” has at least one junction that differs from every annotated transcript. A “new 3′ end” has the same junction chain as an annotated transcript but a 3′ end more than 25 nt away. A “new locus” overlaps no annotated gene.

**Figure 2.**
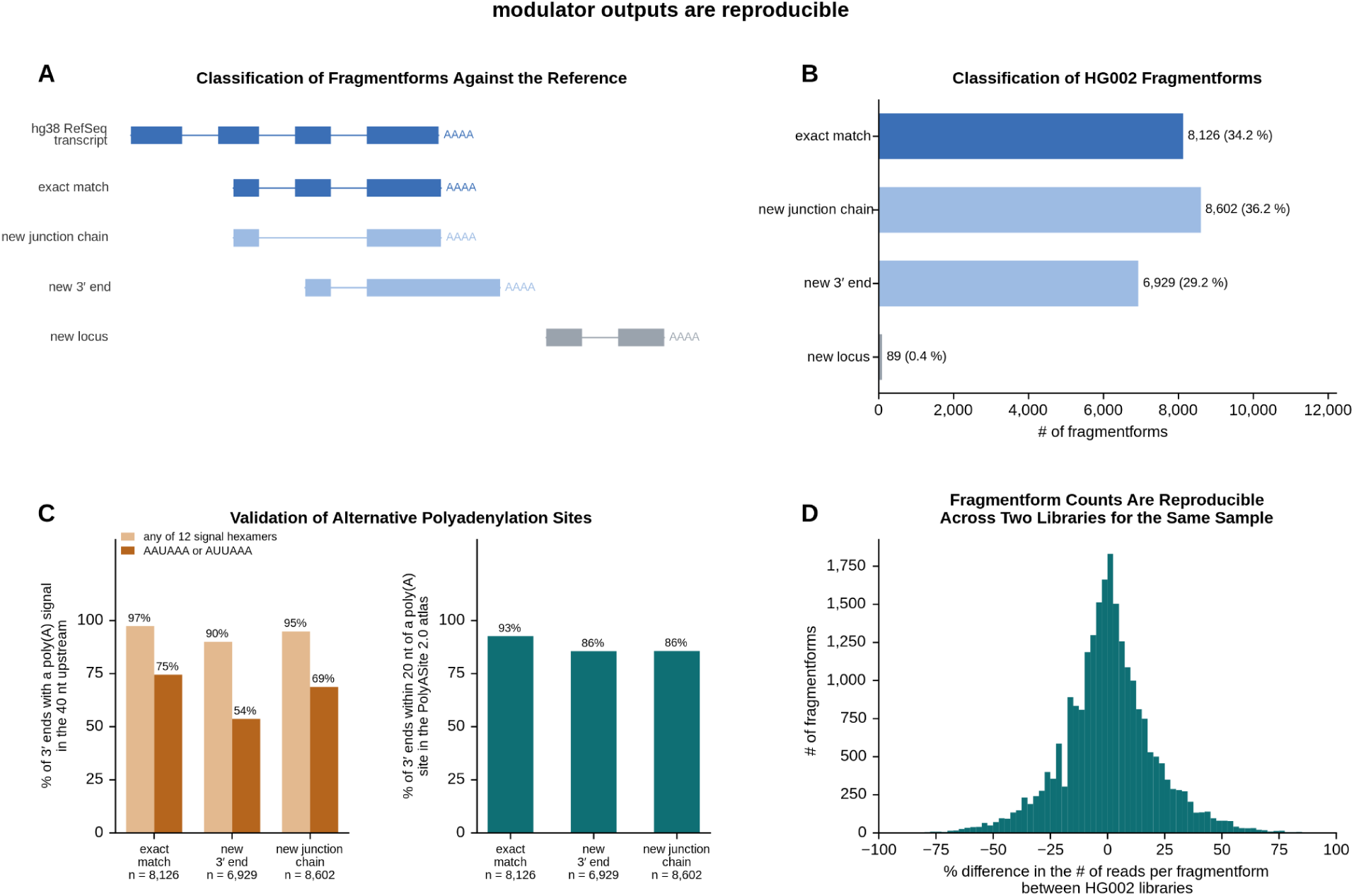
Fragmentforms represent reproducible transcript structures. **(a)** *Classification of fragmentforms against the reference*. Schematic. The blue track is an hg38 RefSeq transcript. Below it are the four classes of fragmentform: an exact match (blue), a new junction chain (light blue), a new 3′ end (light blue), and a new locus (grey). Every fragmentform starts short of the reference 5′ end to demonstrate that the 5′ end does not need to match. An exact match has the same junction chain and a 3′ end within 25 nt of the hg38 RefSeq transcript. A new junction chain has a junction that differs from the hg38 RefSeq transcript. A new 3′ end has the same junction chain and a 3′ end more than 25 nt away from the hg38 RefSeq transcript. A new locus has by default at least two exons and overlaps no annotated gene within the hg38 RefSeq transcriptome. **(b)** *Two thirds of HG002 fragmentforms are absent from the reference transcriptome*. Number of fragmentforms in each class (Figure 2a) from the two HG002 RNA libraries. Of 23,746 fragmentforms across 9,318 genes, 8,126 (34.2%) are exact matches, 8,602 (36.2%) have a new junction chain, 6,929 (29.2%) have a new 3′ end, and 89 (0.4%) are new loci. **(c)** *Validation of alternative polyadenylation sites*. (left) We searched the 40 nt upstream of every fragmentform 3′ end for a poly(A) signal. The percentage of 3′ ends with any of the 12 poly(A) signals (light amber) or with AAUAAA or AUUAAA (dark amber) is shown by class. (right) the percentage of 3′ ends within 20 nt of a PolyASite 2.0 site, by class. **(d)** *Fragmentform counts are reproducible across two libraries for the same sample*. HG002 RNA was sequenced twice on two different flow cells (‘library 1’ and ‘library 2’). For each of the 23,746 fragmentforms, the reads assigned to each sequencing HG002 library were scaled to reads per million, and the percent difference between libraries is plotted as a histogram (dark teal). The median difference is +0.3%, and half of all fragmentforms fall within −10% to +11%. The Spearman correlation of raw counts is 0.96.

From the two HG002 libraries, Modulator built 23,746 fragmentforms in 9,318 genes (**Figure 2b**). Of these, 8,126 (34%) were exact matches and 8,602 (36%) had a new junction chain. Another 6,929 (29%) had a new 3′ end, and 89 were new loci. Thus, two thirds of the fragmentforms did not exactly match a reference transcript by junction chain and 3′ end.

To evaluate the inferred 3′ ends, we tested their support as poly(A) sites, where transcripts are cleaved and poly(A) tails are added. A real poly(A) site is likely to have a poly(A) signal, i.e. a six-nucleotide sequence such as AAUAAA, within 40 nt upstream of it. We also assessed overlap with the PolyASite 2.0 atlas of experimentally determined poly(A) sites (Herrmann *et al*. 2019). We tested both features on every fragmentform 3′ end (**Figure 2c**). One of the 12 known poly(A) signals lay upstream of 97% of exact-match ends, 90% of new 3′ ends, and 95% of new-junction-chain ends, compared to 13% to 17% at shuffled positions (**see Methods**). A PolyASite 2.0 site lay within 20 nt of 93%, 86%, and 86% of the ends, compared to 5% at shuffled positions. These results support the inferred 3′ ends, including those absent from the reference annotation.

We next assessed the reproducibility of fragmentform abundance estimates between the two HG002 libraries. We counted the reads assigned to each fragmentform in each library, scaled the counts to reads per million, and computed the percent difference between libraries (**Figure 2d**). The differences centered on zero, with a median of +0.3%. Half of all fragmentforms fell within −10% to +11%, and the raw counts had a Spearman correlation of 0.96. Thus, fragmentform abundance estimates were reproducible between libraries of the same sample.

### A framework for transcript-, allele-, and condition-associated comparisons

We organized modification analysis around three comparisons: between fragmentforms, between alleles, and between conditions (**Figure 3a**). These comparisons use the transcript-architecture, sequence-variation, and modification information available from individual reads. We then annotated differences between fragmentforms or alleles and changes between conditions across the full data set.

**Figure 3.**
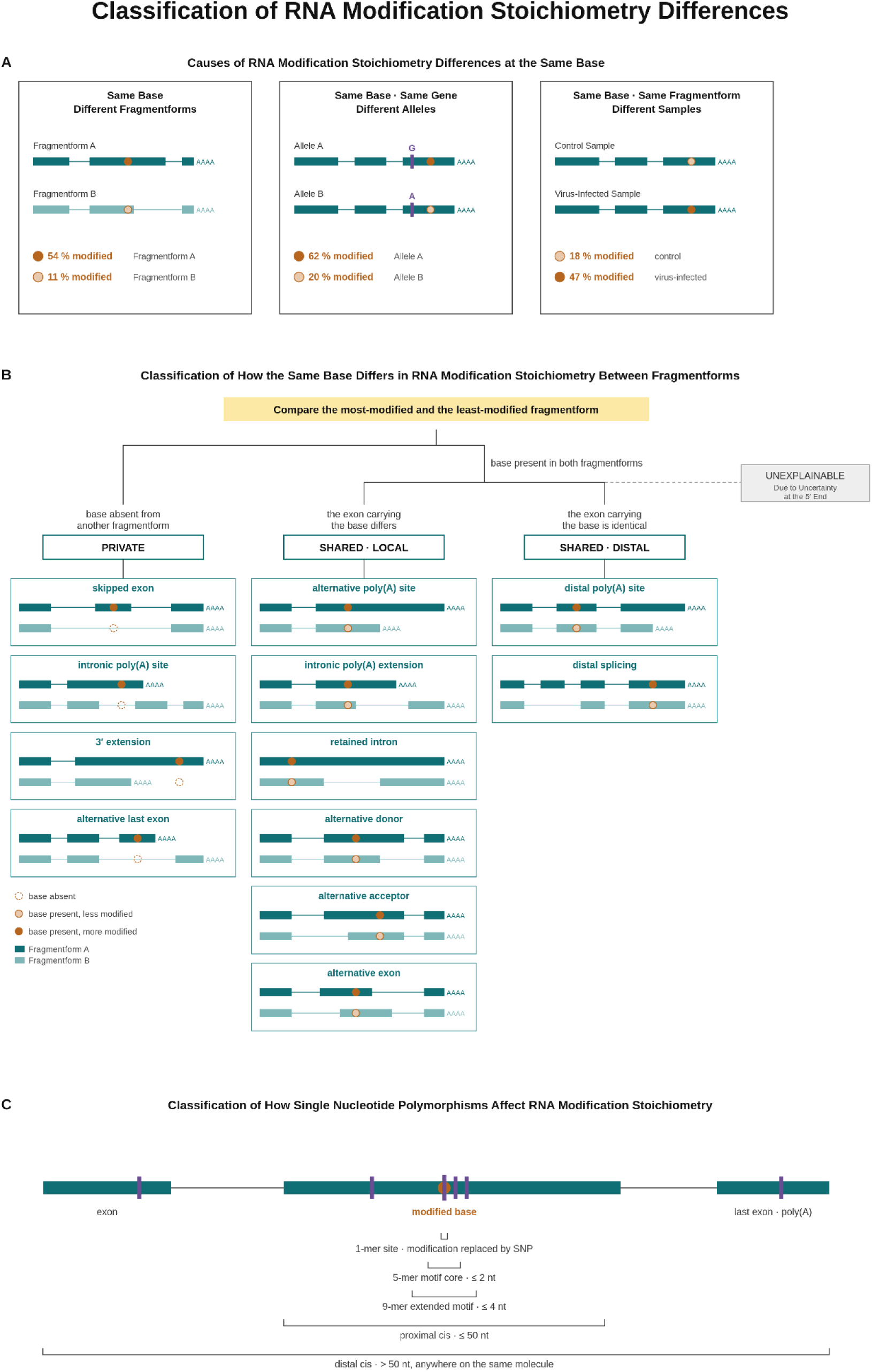
Classification of RNA modification stoichiometry differences. **(a)** *Causes of RNA modification stoichiometry differences at the same site*. Schematic. Three boxes show the same site (amber) that differs between (left) two fragmentforms of one gene, (middle) two alleles of one gene, and (right) one fragmentform but two different samples (e.g. a control and a virus-infected sample). The purple bar marks the SNP. The percentages are example stoichiometries. A filled circle marks the more modified case and an open circle the less modified case. **(b)** *Classification of how the same site differs in RNA modification stoichiometry between fragmentforms*. Schematic. Modulator compares the most modified and the least modified fragmentform at each site. Fragmentform A is dark teal, Fragmentform B is light teal, and the RNA modification site is amber. Sites can be classified as private (left), where the site is absent from another fragmentform. These are sub-classified into their root cause, where the site is absent from another fragmentform due to a skipped exon, an intronic poly(A) site, a 3′ extension, or an alternative last exon. Sites can be classified as “shared local” (middle), where the site is present in both fragmentforms but the two exons where the site is located differ. These are then sub-classified based on the difference between the two exons: an alternative poly(A) site, an intronic poly(A) extension, a retained intron, an alternative donor, an alternative acceptor, and an alternative exon entirely. Lastly, sites can be classified as “shared distal” (right), where the site is present in both fragmentforms, the exon containing the site is identical between the two fragmentforms, and the only difference between the fragmentforms is located elsewhere within the RNA molecules. These are sub-classified into two categories: a distal poly(A) site and distal splicing difference. A dotted circle marks a site absent from that fragmentform. Differences that only a lost 5′ end could explain are set aside as unexplainable. All modification stoichiometries in the figure are representative examples. **(c)** *Classification of how single nucleotide polymorphisms affect RNA modification stoichiometry.* Schematic. A fragmentform is drawn as teal exons with the modified site in amber and SNPs in purple. Each SNP is placed in a distance class according to its distance from the modified site along the spliced RNA. At the nucleotide itself, the modification is replaced by the SNP. Within 2 nt is the motif core (5mer), within 4 nt is the extended motif (9mer), within 50 nt is proximal, and beyond 50 nt anywhere on the same molecule is distal.

To better understand differences in modification stoichiometries between transcripts, we defined a classification hierarchy of the differences between the fragmentforms of one gene (**Figure 3b**). Modulator compares the most modified and the least modified fragmentform at each site. The site may be absent from one fragmentform, because that fragmentform splices it out or ends before it. We refer to such a site present uniquely within the other fragmentform as “private.” These sites are subclassified based on whether they are contained within a skipped exon, an intronic poly(A) site, a 3′ extension, or an alternative last exon, as shown in **Figure 3b**.

Instead of being a private site, the site may be present in both fragmentforms, which we refer to as a “shared” site. Each transcript-associated difference is first classified as either private or shared. Shared sites are then classified according to whether the two fragmentforms differ locally at the site itself or only distally elsewhere in the transcript. When the exon that contains the site differs between the fragmentforms, the site is sub-classified according to the local difference in transcript architecture. These classifications include an alternative poly(A) site, an intronic poly(A) extension, a retained intron, an alternative donor, an alternative acceptor, and an alternative exon, as drawn in **Figure 3b**. A donor is the 5′ boundary of an intron and an acceptor is its 3′ boundary. When the exon that contains the site is identical, the two fragmentforms differ only at a distant poly(A) site or a distant junction. Finally, a difference that depends on the 5′ end is set aside as “unexplainable.” In summary, every difference between fragmentforms receives one of twelve classifications, or is set aside as unexplainable.

To classify allele-associated modification differences, we next defined a second framework based on the distance from the SNP to the modified site (**Figure 3c**). Each significant RNA modification stoichiometry difference associated with a SNP is placed in a distance class by its distance to the site along the spliced RNA. A SNP at the modified nucleotide itself replaces the modification. A SNP within 2 nt sits in the “motif core,” the two nucleotides on either side of the modified nucleotide. A SNP within 4 nt sits in the “extended motif.” A SNP within 50 nt is “proximal,” and a SNP beyond 50 nt on the same molecule is “distal.” In summary, we have developed a distance-based classification system for how SNPs are associated with RNA modification stoichiometries.

To identify condition-dependent changes, Modulator compares modification stoichiometry within the same fragmentform across conditions. Together, these transcript-, allele-, and condition-associated classifications provide the framework used in the following analyses.

### Transcript architecture is associated with RNA modification differences

#### Number of RNA modifications differing between fragmentforms in HG002

We first counted the RNA modification sites that differ in stoichiometry between the fragmentforms of a gene in HG002. Modulator tested 141,621 sites with at least 20 reads in two or more fragmentforms of a gene. We defined a site as differing at a false discovery rate of 5% or less and with a stoichiometry difference of at least 10 percentage points. In total, 3,446 RNA modification sites differed between fragmentforms: 3,214 m6A, 177 inosine, 25 pseU, 25 m5C, and 5 2′OmeG (labeled Gm in **Figure 4**) sites. Of these, 3,413 are drawn in **Figure 4**. The remaining 33 are excluded since they involve comparisons with soft-clipped bases. The alternative poly(A) site class held the most sites, 1,951, followed by retained introns with 450 and distal splicing with 335. Only 106 differences were unexplainable. These results demonstrate that the fragmentform assigns an informative class to 97% of the sites that differ between the fragmentforms of a gene, and that the largest class is the alternative poly(A) site.

**Figure 4.**
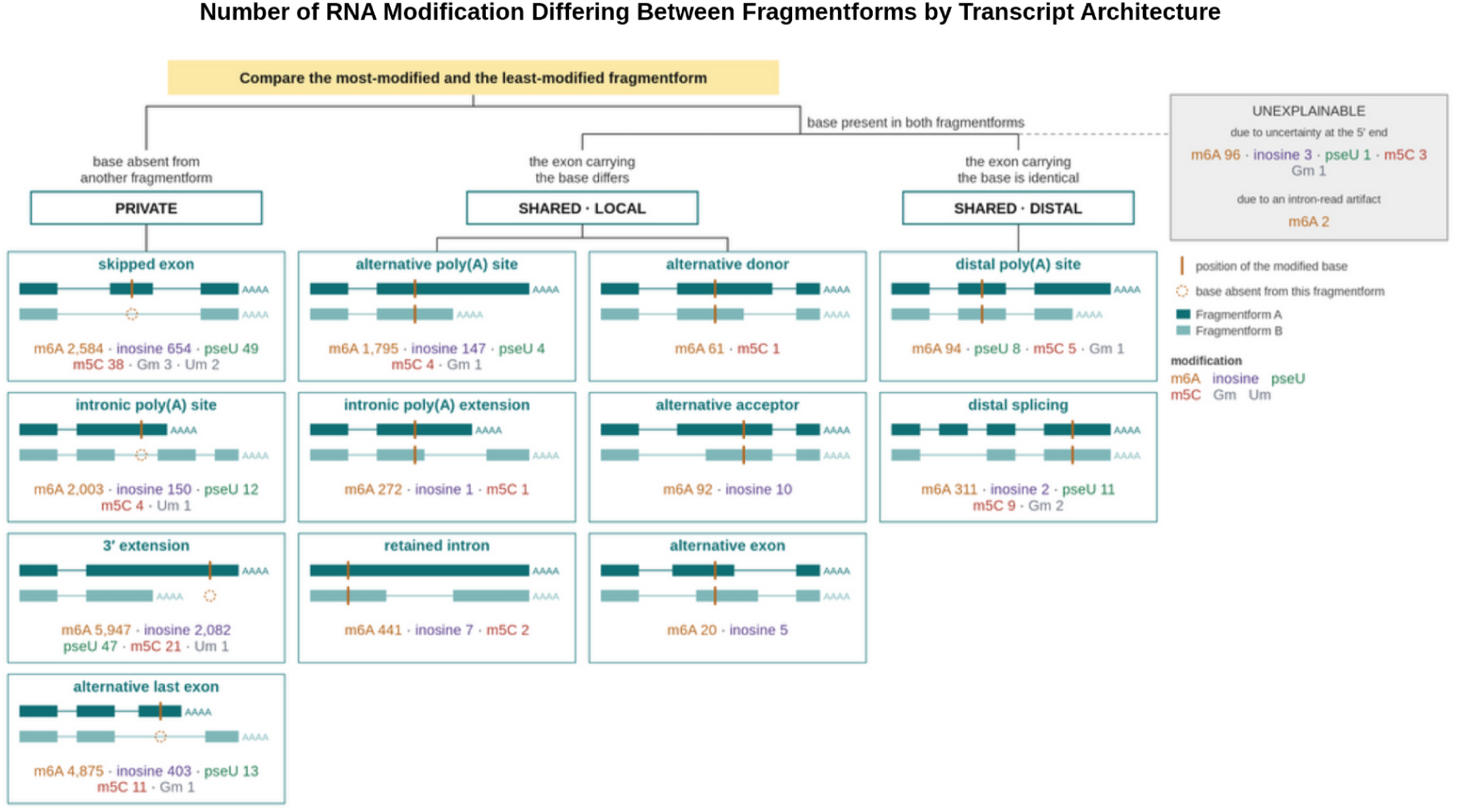
Number of RNA modifications differing between fragmentforms in HG002. After running Modulator on the HG002 direct-RNA sequencing data, we report the number of RNA modification sites by modification (m6A, inosine, pseU, m5C, Gm, Um) that fall into each classification for how the same site differs in RNA modification stoichiometry between fragmentforms (Figure 3b). An amber tick marks the site, and a dotted circle marks a site absent from that fragmentform. There are 3,413 total RNA modification differences between fragmentforms where the site is present in both. There are 18,901 sites that are private to a single fragmentform.

We next counted how many RNA modification sites are private to one fragmentform. Modulator found 18,901 private sites (**Figure 4**). Of these, 8,098 lay in a 3′ extension, 5,303 in an alternative last exon, 3,330 in a skipped exon, and 2,170 at an intronic poly(A) site. By modification, 15,409 were m6A and 3,289 were inosine. These private sites capture differences in the presence of modified sequence, distinct from stoichiometry differences at sites shared between fragmentforms.

#### m6A differences between fragmentforms are associated with distance to nearby junctions and poly(A) sites

For each shared-site class, we next calculated which fragmentform had higher modification stoichiometry. We focused on m6A because it accounted for most shared-site stoichiometry differences. At an alternative poly(A) site, the fragmentform with the distal poly(A) site was more modified at 1,596 sites and less modified at 199 (**Figure 5a**). At an intronic poly(A) extension, the fragmentform that extends to the intronic poly(A) site was more modified at 271 sites and less modified at 1 (**Figure 5b**). Retained introns were more balanced, with 237 versus 204 sites (**Figure 5c**). At an alternative donor or acceptor, the fragmentform with the longer exon was more modified at 54 of 61 and 59 of 92 sites, respectively (**Figure 5d, e**); we observed few alternative exon sites (**Figure 5f**). Distal poly(A) comparisons were relatively balanced (**Figure 5g**). Across several classes, higher m6A stoichiometry was associated with the fragmentform retaining more RNA sequence, although this pattern was not universal.

**Figure 5.**
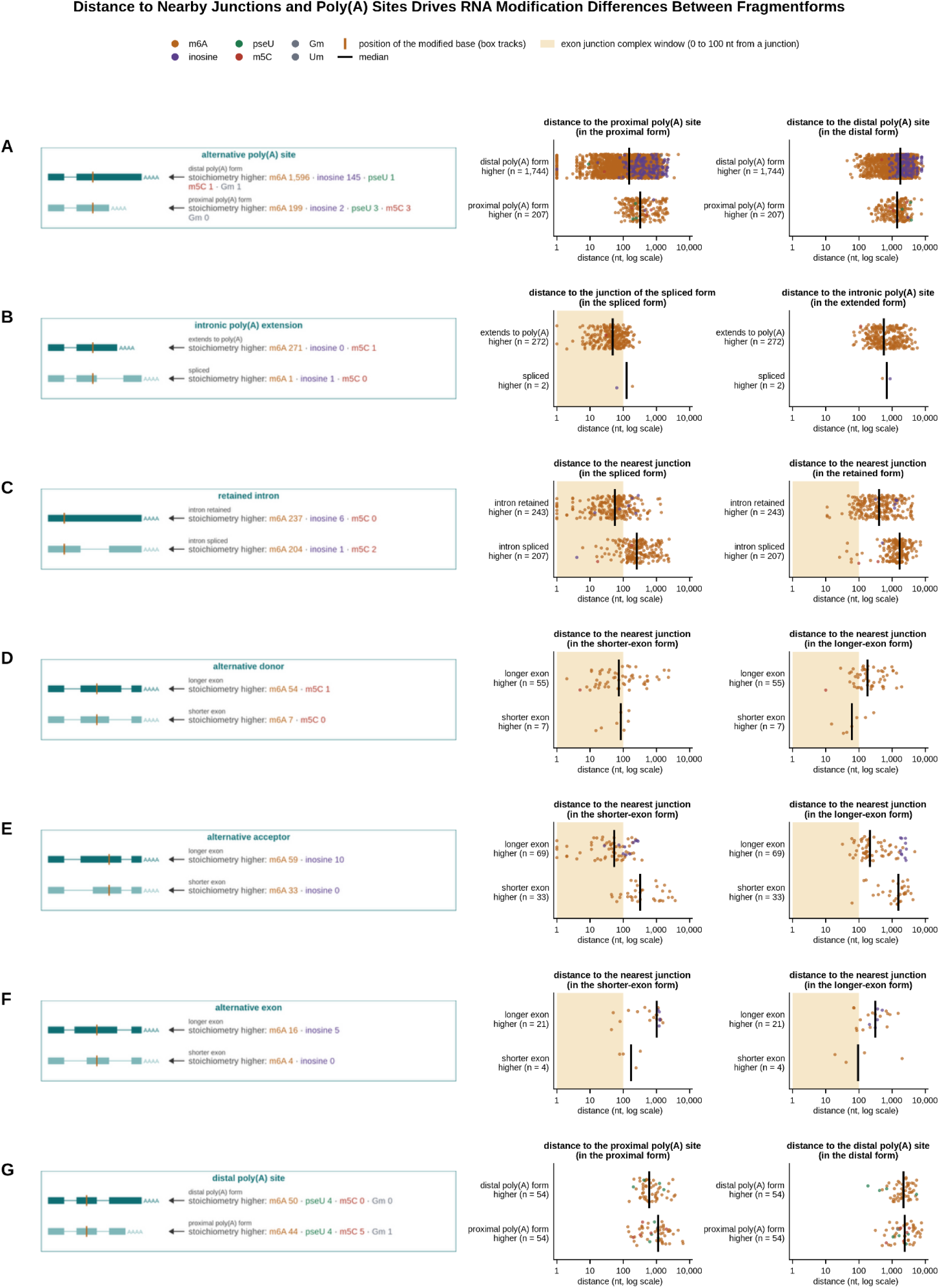
Distance to nearby junctions and poly(A) sites drives RNA modification differences between fragmentforms. Based on the results from Figure 4, the “shared-local” and “shared-distal” classes are further subdivided based on which “form” of the transcript has a higher RNA modification stoichiometry. The plots on the right show RNA modification sites (dots) based on their distance to either splice junctions or poly(A) sites. The number of sites is labeled as the “n” number and the black tick is the median distance. The yellow shading represents any RNA modification sites that are 0 to 100 nucleotides (nt) from a splice junction, demonstrating reported suppression by the exon junction complex. The x axis for distance is logarithmic. **(a)** *Alternative poly(A) site.* Distance to the proximal and to the distal poly(A) site. For m6A, the distal poly(A) site fragmentform is more modified at 1,596 sites and the proximal at 199, with medians of 129 and 326 nt from the proximal site. For inosine, 145 are more modified in the distal poly(A) site fragmentform compared to 2 in the proximal poly(A) site fragmentform. **(b)** *Intronic poly(A) extension.* Distance to the junction of the spliced fragmentform and to the intronic poly(A) site. The intronic poly(A) site fragmentform is more modified at 271 m6A sites compared to the spliced fragmentform at 1 m6A site, with medians of 48 and 189 nt from the splice junction, respectively. **(c)** *Retained intron.* Distance to the nearest junction in the spliced and in the retained fragmentform. The retained fragmentform is more modified at 237 m6A sites and the spliced at 204, with medians of 55 and 258 nt from the junction in the spliced fragmentform. **(d)** *Alternative donor.* Distance to the nearest junction in the shorter-exon and in the longer-exon fragmentform. The longer exon is more modified at 54 m6A sites and the shorter at 7 m6A sites. The median distance to the shorter splice junction is similar for both groups (84 and 85 nt). **(e)** *Alternative acceptor.* Distance to the nearest junction in the shorter-exon and in the longer-exon fragmentform. The longer exon is more modified at 59 m6A sites and the shorter at 33, with median distance of 47 nt and 325 nt from the junction in the shorter-exon fragmentform. For inosine, 10 sites are more modified in the longer-exon fragmentform compared to 0 sites in the shorter-exon fragmentform. **(f)** *Alternative exon.* Distance to the nearest junction in each fragmentform. The longer exon is more modified at 16 m6A sites and the shorter at 4. **(g)** *Distal poly(A) site.* Distance to the proximal and to the distal poly(A) site. The distal fragmentform is more modified at 50 m6A sites and the proximal at 44. The sites in both groups lie 600 to 1,300 nt from the proximal site.

We therefore asked whether the distance from the site to the separating feature explains which fragmentform is more modified. The separating feature is either the junction or poly(A) site that one fragmentform uses and the other does not. We measured the distance along the spliced RNA from every differing site to this feature (**Figure 5a to g, right**). At intronic poly(A) extensions, the sites where the extended fragmentform was more modified sat a median of 48 nt from the junction, and 83% of sites sat within 100 nt. At retained introns, the sites where the retained fragmentform was more modified sat 55 nt from the junction, whereas the sites where the spliced fragmentform was more modified sat 258 nt away. At alternative acceptors, the two groups sat 47 and 325 nt from the junction. At alternative donors, the two groups did not separate (84 and 85 nt). These results demonstrate that an m6A site within about 100 nt of a splice junction (the yellow shading in **Figure 5**) is generally less modified in the fragmentform that splices at that junction.

We observed a similar association near poly(A) sites: shared m6A sites within about 1 kb of a proximal poly(A) site were generally ∼24 percentage points less modified in the fragmentform that used that site than in a fragmentform that extended to a distal site. Of 29,191 tested m6A comparisons at alternative poly(A) sites, 1,795 differed. Of the 1,795 m6A sites within alternative poly(A) fragmentforms, 729 lay within 100 nt of the proximal poly(A) site, where the distal fragmentform was more modified at 717 sites compared with only 12 sites in which the proximal fragmentform was more modified. The excess of distal-higher sites persisted from 100 to 1,000 nt (794 compared to 145), and was substantially reduced beyond 1 kb (85 compared to 42).

We finally asked whether inosine follows the same distance dependence as m6A. At alternative poly(A) sites, the distal fragmentform was more edited at 145 sites and less edited at 2. These sites sat a median of 420 nt from the proximal site. While inosine also showed a strong distal-fragmentform bias, the relationship to distance appeared distinct from that observed for m6A.

### Modulator detects allele-associated RNA modification differences

Before examining allele-associated modification differences, we compared RNA-derived SNP calls with the HG002 Genome in a Bottle benchmark (Zook *et al*. 2019, Wagner *et al*. 2022, Hansen *et al*. 2026). Modulator called 14,759 SNPs from the two HG002 libraries. Of the 13,576 calls in the regions where the benchmark is considered reliable, 12,640 (93%) were true heterozygous variants, i.e. positions where the two copies of the genome differ (**Figure 6a**). Precision rose with read depth, from 75% at 20 to 50 reads to 98% above 200 reads. Overall, RNA-derived calls achieved 93% precision within high-confidence benchmark regions and recovered 93% of benchmark variants with at least 20 reads in the scanned exons.

**Figure 6.**
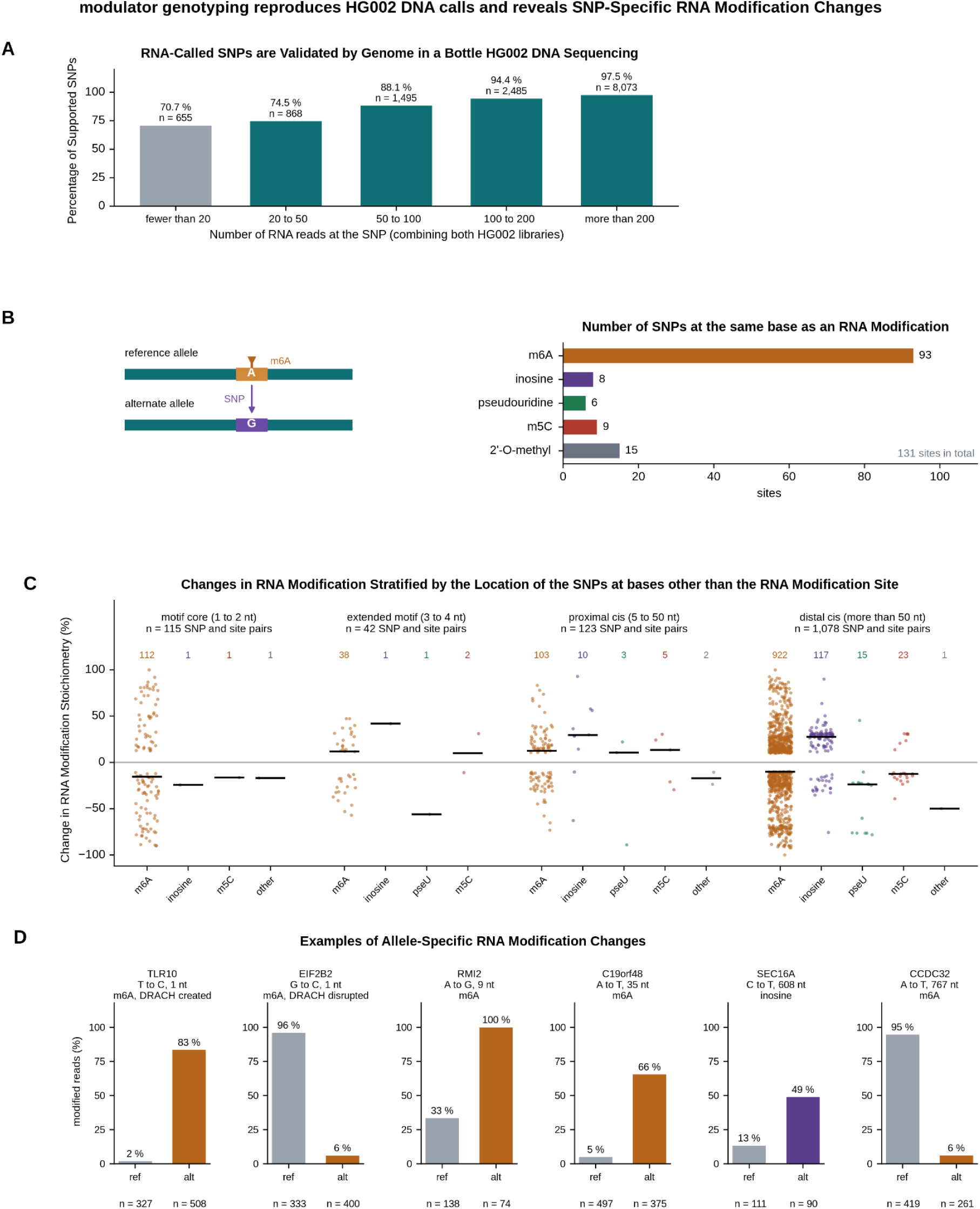
Modulator genotyping reproduces HG002 DNA calls and reveals SNP-specific RNA modification changes. **(a)** *RNA-called SNPs are validated by Genome in a Bottle HG002 DNA sequencing*. Modulator called 14,759 SNPs from the two HG002 RNA sequencing samples. Of these, 13,576 SNPs fall within the high-confidence regions of the Genome in a Bottle benchmark. Bars (dark teal) show the percentage of calls in the high-confidence regions of the benchmark that are true heterozygous variants, by the number of RNA reads at the SNP: 70.7% below 20 reads (n = 655, grey because the allele tests require at least 20 reads), 74.5% at 20 to 50 reads (n = 868), 88.1% at 50 to 100 reads (n = 1,495), 94.4% at 100 to 200 (n = 2,485), and 97.5% above 200 (n = 8,073). Within the high-confidence regions, Modulator called 93.0% of the heterozygous variants it tested with at least 20 reads. The Modulator haplotyping feature recorded no haplotyping assignment errors within 3,724 phased SNP haplotypes when examined against the Genome in a Bottle trio-phased benchmark. **(b)** *131 SNPs replace the modified nucleotide itself*. A SNP occurring at the same position as an RNA modification results in loss of the RNA modification or the opportunity for a new RNA modification to arise. The schematic on the left shows a reference allele whose adenosine is m6A-modified and an alternate allele in which a SNP has replaced that adenosine with a guanosine. This demonstrates how allele variation between homologous chromosomes can result in differential RNA modification. We observed 93 such sites for m6A, 8 sites for inosine, 6 sites for pseU, 9 sites for m5C, and 15 sites for the 2′-O-methyl modifications. **(c)** *Most allele effects act from more than 50 nt away*. Each dot is an RNA modification site that is altered by a SNP. The y-axis is calculated by taking the alternate allele stoichiometry minus the reference (hg38) allele stoichiometry. The RNA modification sites are distributed across four plots based on the distance of SNP to the RNA modification site: (left) the motif core at 1 to 2 nt (n = 115), (left, middle) the extended motif at 3 to 4 nt (n = 42), (middle, right) proximal at 5 to 50 nt (n = 123), and (right) distal at more than 50 nt (n = 1,078). The count per modification is printed above and the black tick is the median RNA modification difference. The median absolute effect is 47 percentage points in the motif core and 23 to 27 for the other distances. In the motif core, a SNP that creates a DRACH motif raises the m6A on that allele at 28 of 29 sites, and a SNP that destroys one lowers it at 35 of 35. **(d)** *Examples of allele-specific RNA modification changes*. For six RNA modification sites, the stoichiometry on the reference allele (grey) and the alternate allele (modification color) are shown. Reads are pooled over the two HG002 direct-RNA sequencing samples with read counts (n) under each bar. The site-level statistics are as follows: TLR10, T to C at 1 nt, DRACH created, m6A 2% to 83% (327 and 508 reads). EIF2B2, G to C at 1 nt, DRACH disrupted, 96% to 6% (333 and 400). RMI2, A to G at 9 nt, 33% to 100% (138 and 74). C19orf48, A to T at 35 nt, 5% to 66% (497 and 375). SEC16A, C to T at 608 nt, inosine 13% to 49% (111 and 90). CCDC32, A to T at 767 nt, 95% to 6% (419 and 261).

We next asked whether the phase of the SNPs, i.e. which alleles lie on the same molecule, matches the genome. Modulator phases SNPs that lie on the same molecules into groups of 2 to 4 SNPs, and it found 4,402 such groups in HG002. The benchmark assigns each allele to the maternal or paternal copy using the genomes of the parents. Of the 4,402 groups, 3,724 contained at least two SNPs whose phase is known in the benchmark and determined by Modulator. A “switch error” is a pair of neighboring SNPs whose assignment to the two alleles disagrees with the benchmark. The 6,858 adjacent SNP pairs in the 3,724 evaluable groups had 0 switch errors. These results demonstrate that the SNPs and their phase read from RNA match this genome without any phasing errors.

We next examined how genetic variation influences RNA modification stoichiometry. The most direct mechanism occurs when a SNP replaces a modified nucleotide. For instance, an A-to-G SNP at an m6A site removes the adenosine required for modification on one allele, whereas a G-to-A SNP creates a potentially modifiable adenosine on the other allele. Within HG002, Modulator found 131 such sites (**Figure 6b**): 93 m6A sites, 15 2′-O-methyl sites, 9 m5C sites, 8 inosine sites, and 6 pseU sites. However, these direct effects accounted for only a small fraction of allele-associated modification differences. Across 314 genes, Modulator identified 1,358 statistically significant RNA modification differences associated with 999 SNPs and 427 RNA modification sites. Of these SNP-site pairs, 115 fell within the motif core, 42 within the extended motif, 123 within 50 nt, and 1,078 more than 50 nt away (**Figure 6c**). Although motif-core variants had the largest effects, with a median difference of 47 percentage points compared with 23 to 27 in the other distance classes, most allele-associated differences involved SNPs located far from the modified site. These results indicate that most allele-associated RNA modification differences cannot be explained by direct disruption of the modification motif.

We next examined whether motif-core variants behaved as expected from known m6A sequence preferences. The m6A writer complex preferentially methylates the DRACH motif (D = A, G, U; R = A, G; H = A, C, U) (Meyer *et al*. 2012, Linder *et al*. 2015). We found that a SNP that created a DRACH motif raised the m6A on that allele at 28 of 29 sites, whereas a SNP that disrupted a DRACH motif lowered the m6A on that allele at 35 of 35 sites. These results indicate that motif-core variants affect m6A in the expected direction, even though most allele-associated modification differences involve SNPs located outside the canonical motif.

We additionally highlight representative examples of large allele-associated RNA modification differences across the full range of SNP distances (**Figure 6d**). These examples illustrate that a single nucleotide change can likely lead to increases in modification stoichiometry from 2% to 83% or decreases from 96% to 6%, regardless of whether the associated SNP lies within the canonical motif or hundreds of nucleotides away.

### Zika virus infection changes RNA modification stoichiometry within the same fragmentforms

We next asked whether RNA modification stoichiometry changes between conditions within a fixed fragmentform context. We applied Modulator to DRS data from three mock-infected and three ZIKV-infected Huh7 samples that we previously generated (Aufgebauer *et al*. 2026), one library per sample, six flow cells in total. Modulator detected significant changes within the same fragmentform at 1,253 fragmentform and site pairs across five modifications (**Figure 7a**): 270 m6A, 439 inosine, 471 pseU, and 72 m5C sites, plus one 2′OmeU site not shown. These comparisons identify infection-associated modification changes within a defined fragmentform context.

**Figure 7.**
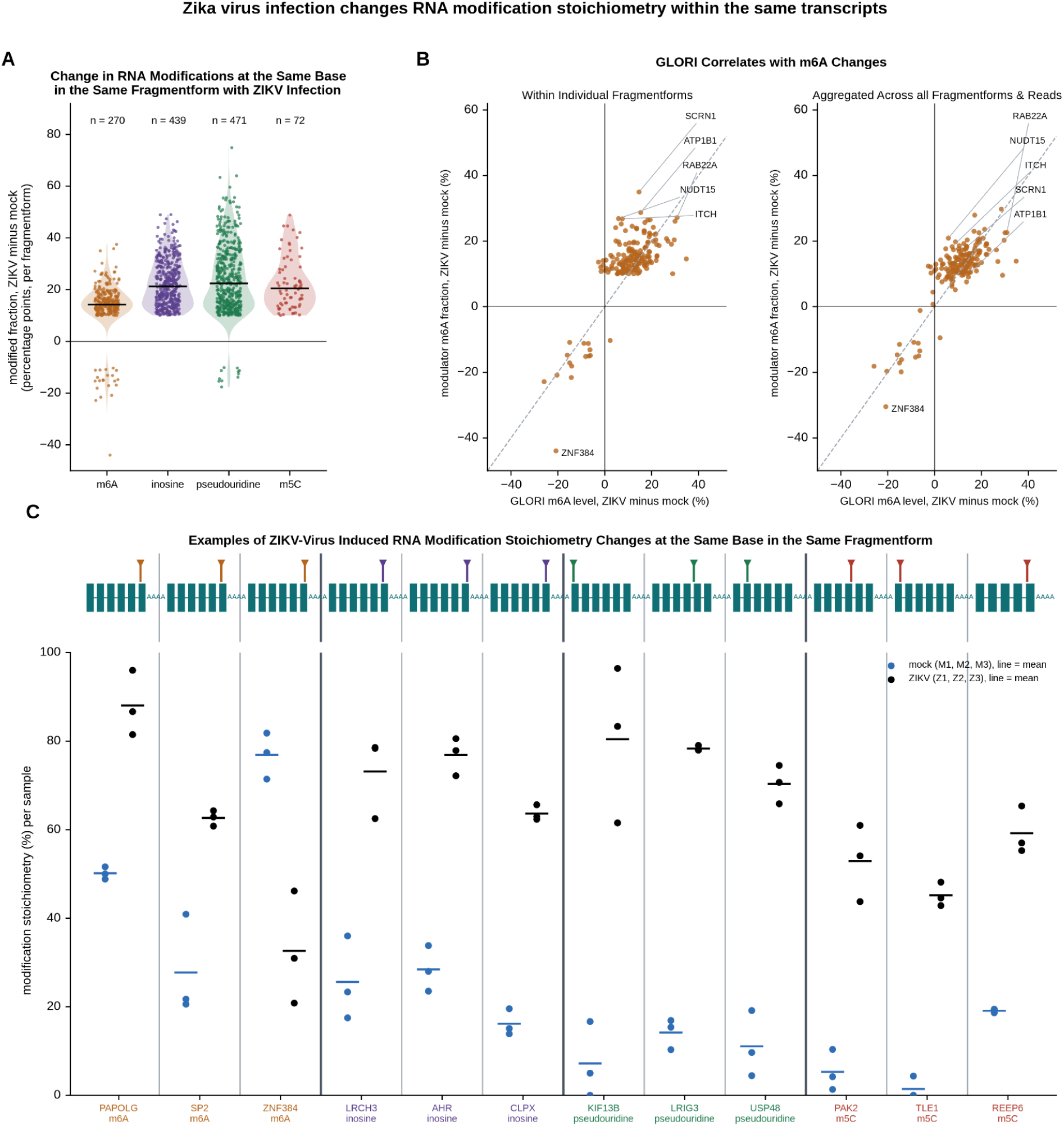
Zika virus infection changes RNA modification stoichiometry within the same transcripts. **(a)** *Infection raises inosine, pseU, and m5C site stoichiometry in the same fragmentform*. Each dot is a RNA modification site whose stoichiometry differs between mock and ZIKV-infected Huh7 cells. The y-axis is calculated based on the RNA modification stoichiometry of the ZIKV-infected condition minus the RNA modification stoichiometry of the Mock condition. The violin shows the distribution and the black tick is the median modification stoichiometry difference. There exist n = 270 m6A, 439 inosine, 471 pseU, and 72 m5C differential RNA modification sites. The median stoichiometry differences are +14, +21, +22, and +20 percentage points, respectively. **(b)** *GLORI correlates with m6A changes*. GLORI, a chemical method that measures m6A at single nucleotides, was run on the same six libraries. For differential m6A RNA modification sites in single fragmentforms (n=270), there were 183 sites that were also measured by GLORI. (left plot) Of these 183 sites, 176 change in the same direction, with a Spearman correlation of 0.56 when correlated with the m6A levels on the individual fragmentform associated with the differential m6A RNA modification. (right plot) The correlation with GLORI improves with a site-level aggregation at the same sites since GLORI does not measure transcript level modifications (Spearman correlation of 0.65). **(c)** *Examples of virus-specific RNA modification changes*. We plot the three largest changes in RNA modification stoichiometry for each of the four modifications (m6A, inosine, pseU, m5C) where virus-specific RNA modifications were identified. The mock samples are plotted in blue and the infected samples in black. The corresponding lines are the condition’s mean RNA modification stoichiometry. Above each example, the fragmentform is drawn with its 3′-most exons in teal, the poly(A) tail as AAAA, and the modified site marked in the modification color. Reads per library at the site range from 18 to 987 and are listed in **Supplementary Table 2.**

Most condition-associated changes were increases in modification stoichiometry. Remarkably, inosine, pseU, and m5C increased at 971 of 982 sites, whereas m6A increased at 249 sites and decreased at 21 sites (**Figure 7a**). This directional pattern suggests that infection is associated predominantly with increased modification stoichiometry across multiple modifications.

We compared the Modulator-detected m6A changes with existing GLORI measurements from the corresponding samples reported by (Aufgebauer *et al*. 2026). GLORI measures m6A stoichiometry at single-nucleotide resolution using an independent chemical approach (Liu *et al*. 2023, Shen *et al*. 2024, Sun *et al*. 2025). We compared the change between conditions at every m6A site that both methods could test. At the 183 sites with a significant Modulator change and a GLORI measurement, 176 (96%) changed in the same direction, with a Spearman correlation of 0.56 (**Figure 7b, left**). GLORI does not measure transcript-level modification, so we also pooled the Modulator reads across all fragmentforms at the same sites. At the 173 sites with a pooled measurement, 166 (96%) agreed, and the Spearman correlation rose to 0.65 (**Figure 7b, right**). The reason that ten sites were lost is because the same m6A site can be changed with virus infection across multiple fragmentforms which were aggregated into a single pooled measurement. These comparisons support the direction of infection-associated m6A changes at overlapping genomic positions, but do not independently establish their transcript specificity.

Replicate-level examples illustrate the three largest changes for each of four modifications (m6A, inosine, pseU, m5C; **Figure 7c**). In PAPOLG, m6A rose from 52%, 50%, and 49% in mock-infected samples to 82%, 87%, and 96% in infected samples. In KIF13B, pseU rose from 0%, 5%, and 17% to 83%, 96%, and 62%. In TLE1, m5C rose from 4%, 0%, and 0% to 43%, 45%, and 48%. Together, these results show that Modulator detects condition-dependent modification changes within a defined fragmentform context, with orthogonal support from GLORI for the direction of m6A changes at overlapping positions.

## DISCUSSION

### Summary

RNA modifications are deposited on individual molecules, yet they are usually reported at aggregated genomic positions, which discards the transcript context in which each molecule was modified. Preserving that context is the central challenge for transcript-level analysis of RNA modifications with Nanopore direct-RNA sequencing, because most reads lack the 5′ end needed to assign them to an annotated transcript. We introduce “fragmentforms” in Modulator to retain the transcript context that every included read reports, namely its 3′ end and junction chain, thereby recovering transcript-level RNA modification information from Nanopore direct-RNA sequencing data. For the HG002 transcriptome, Modulator built 23,746 fragmentforms, two thirds of which contain transcript variants absent from the reference transcriptome. These features would not have been captured by aligning to the hg38 reference transcriptome alone, and underscore the importance of constructing sample-specific reference transcriptomes directly from genomic alignments.

Including 5′-truncated reads substantially increased the number of fragmentforms and of testable and differential RNA modification sites. This gain reflects both the additional read depth contributed by retained reads and the ability to preserve their observable transcript context, enabling comparisons that would otherwise not be possible. We were therefore able to make more confident transcriptome-wide observations of RNA modification biology within HG002. These observations reveal differences associated with the transcript architecture, genotype, or condition, illustrating the value of preserving transcript context. The transcript-architecture findings most directly demonstrate the value of the approach because they emerge only when modification measurements are interpreted in the context of transcript architecture rather than from site-level measurements.

Firstly, we find that transcript architecture is strongly associated with m6A stoichiometry. In particular, modification differences between fragmentforms depend on the distance of a site to nearby splice junctions and poly(A) sites. We observed that an m6A site within about 100 nt of a splice junction is less modified in the spliced transcript. This pattern is consistent with the previously described depletion of m6A near splice junctions and exon-intron boundaries (Yang *et al*. 2022, He *et al*. 2023, Luo *et al*. 2023, Uzonyi *et al*. 2023), which may in part reflect exclusion by the exon junction complex.

Our transcript-resolved analysis further indicates that use of a proximal poly(A) site is associated with lower m6A stoichiometry at nearby shared sites than use of a distal poly(A) site. This pattern occurs in a transcript region with well-established m6A enrichment and is compatible with evidence that transcriptional dynamics influence regional m6A deposition (Meyer *et al*. 2012, Wang *et al*. 2024). Our companion study, which also used Modulator to resolve isoform-specific m6A, identified proximal poly(A) sites as positive determinants of m6A accumulation and defined cleavage-dependent and cleavage-independent mechanisms through which the polyadenylation machinery reshapes m6A deposition (Aufgebauer et al. 2026). The two studies therefore address different levels of the same phenomenon. The companion study identifies mechanisms linking poly(A)-site architecture to m6A deposition, whereas the present study reveals that poly(A)-site usage is associated with systematic differences in m6A stoichiometry across thousands of transcript comparisons. Together, the two studies identify poly(A)-site architecture as a major determinant of the m6A landscape.

The exact mechanism underlying every poly(A)-site-associated difference observed here is not established. Our companion study provides evidence that cleavage-dependent and cleavage-independent functions of the polyadenylation machinery can reshape m6A deposition in specific transcript contexts (Aufgebauer et al. 2026). However, the transcriptome-wide associations observed in the present study suggest that additional mechanisms are also likely to contribute. These could include additional effects on m6A writer recruitment or access, differences in RNA processing kinetics, other non-coding RNAs or enzymes, or feedback between m6A deposition and poly(A)-site selection. Our data here establish a strong association between poly(A)-site usage and local m6A stoichiometry, but they do not determine the mechanism underlying every observed difference.

In comparison to m6A, the distance to either splice junction or poly(A) site does not generally explain differences in inosine stoichiometry between transcripts. The inosine differences at alternative poly(A) sites lie a median of 420 nt from the proximal site, far outside the range over which m6A responds. Inosine is deposited by the ADAR enzymes, which require double-stranded RNA, so the inosine differences between fragmentforms are more likely to depend on which fragmentform contains the sequence that forms the double-stranded partner.

Secondly, transcript-associated modification differences are not explained solely by proximity to splice junctions or poly(A) sites. For instance, we observe 199 m6A sites that are higher within the proximal poly(A) transcript when compared with the distal poly(A) transcript, which accounts for ∼10% of all differential m6A sites in alternative poly(A) fragmentforms. The mechanism for these differences is currently unknown.

Thirdly, and similarly, we compared m6A between spliced transcripts and transcripts retaining their introns. For the transcripts retaining their introns, many of the increased m6A sites are well explained by distance to the nearby splice site. Surprisingly though, ∼50% of comparisons between spliced transcripts and transcripts retaining their intron have increased m6A stoichiometry in the spliced transcript. Possible mechanisms could include differences in transcript-processing state, with unspliced transcripts representing earlier processing intermediates, variations in splicing machinery, or a model in which the mechanisms driving intron retention exclude m6A.

Fourthly, the distal poly(A) site class is similarly split, with 50 sites higher in the distal fragmentform and 44 higher in the proximal fragmentform, and does not show a relationship with distance. These patterns suggest that transcript-associated modification differences can arise through mechanisms beyond proximity to splice junctions or poly(A) sites. Such mechanisms could include differences in RNA structure, RNA processing, or other transcript features that alter site accessibility (Bass 2002, Höfler and Duss 2024, Shachar *et al*. 2024). More broadly, these findings suggest that transcript-resolved modification measurements may provide insight into molecular features of RNA that are not apparent from sequence alone.

Fifthly, SNPs can be associated with RNA modification differences *in cis* even if they are far away from the RNA modification site. SNPs that disrupt or create motifs have predictable effects. Yet, 1,078 of the 1,358 SNP-site pairs involved a SNP more than 50 nt from the modified site. These distal associations suggest that mechanisms beyond direct disruption of the modification motif contribute to allele-associated modification differences. Because our allele analysis pools reads across all fragmentforms of a gene, an apparent allele-associated modification difference may reflect either a direct effect on modification or an allele-dependent difference in fragmentform usage. The current analysis does not distinguish between these possibilities. Other distal associations could reflect RNA structure, binding of regulatory proteins, or combinations of these effects (Su *et al*. 2018, Freund *et al*. 2020, Uzonyi *et al*. 2023, Shachar *et al*. 2024, Park and Cenik 2025). Modulator provides a framework for identifying these associations and prioritizing them for mechanistic investigation.

Finally, ZIKV infection changes RNA modification stoichiometry within the same fragmentform, separate from a change in which transcripts are expressed. Inosine, pseU, and m5C were increased at almost every changed site, while m6A increased at most sites. Because each sample was sequenced as a single library on a single flow cell, technical effects cannot be fully excluded, particularly for the strongly directional increases in inosine, pseU, and m5C. GLORI agreed in direction for 96% of the m6A changes measured by both methods. This agreement supports the direction of m6A changes at overlapping genomic positions but does not independently establish fragmentform specificity. The cause of these increased modification stoichiometries is not known. Nevertheless, these results demonstrate the exciting opportunity that Modulator offers to explore condition-dependent changes in RNA modification stoichiometry within a defined fragmentform context.

### Limitations

We acknowledge several limitations. First, fragmentforms lack definite 5′ ends by design. This means that fragmentforms are not complete transcripts and do not provide full-length isoform reconstruction. They are a construct primarily used to examine differences in RNA modification stoichiometries between observable transcript architectures. Differences that could be explained only by 5′ end features that the reads do not observe are set aside as unexplainable. These made up 3% of the differential sites in HG002. Fragmentforms are intentionally not transcript reconstructions. Unlike transcriptome assembly approaches, fragmentforms do not attempt to determine complete transcript structures. These fragmentforms preserve the transcript context directly observable in each read for modification analysis. As a result, fragmentforms and transcript assemblies address related but distinct analytical goals.

Second, a heavily 5′-truncated read that is compatible with more than one fragmentform is assigned to the most abundant one, so its membership at any feature upstream of where it terminated is inferred rather than observed. At shared sites near the 3′ end of fragmentforms that diverge far upstream, this mixing of molecules can mask a real difference that would be resolved by using only the reads that span the point of divergence, at the cost of read depth. We are actively developing a truncation-aware differential test that identifies such sites by repeating each comparison with only the reads that span the point of divergence.

Third, reads with low mapping quality, reads without a splice junction, and fragmentforms supported by fewer than 40 reads are excluded. Thus, not every mapped read is assigned to a fragmentform; 16.1 of 28.2 million mapped HG002 reads were assigned under the settings used here, but other settings are possible. The allele comparison also pools the reads of all fragmentforms of a gene and is stratified by library but not by fragmentform, so an allele effect can include a difference in which fragmentform each allele produces. The current analysis does not separate allele-associated transcript usage from modification differences within a fixed transcript context. We are actively developing a separate module that identifies SNP-associated differential transcript usage. These results can be overlapped with identified allele-specific modifications to distinguish transcript-architecture-associated differences from modification differences within a fixed transcript context.

Fourth, we were only able to verify the m6A between-sample results independently with GLORI. For the other modifications, we rely on validation statistics published by Oxford Nanopore Technologies (https://epi2me.nanoporetech.com/rna-mod-validation-all5mer-2026.07/). Calls for these non-m6A modifications should therefore be interpreted cautiously, as performance of nanopore modification-detection workflows remains modification-, model-, and dataset-dependent (Wu *et al*. 2026).

### Features under development

Modulator contains several capabilities that are under active development. These include a truncation-aware differential test; analysis of differential poly(A)-tail length; testing modification co-occurrence on the same molecule; haplotype-aware analysis of modification and transcript usage; annotation of mature-mRNA sequence elements and modifications within them; differential usage analysis of fragmentforms, poly(A) sites, and splice junctions; and an HTML report with an interactive gene browser.

## Conclusion

Modulator assigns informative direct-RNA reads to fragmentforms and distinguishes transcript-associated, allele-associated, and condition-associated differences in RNA modification. Because Modulator does not require a reference transcriptome for fragmentform construction, it may be useful when annotation is incomplete, including in specific cell types, disease states, and non-model organisms, although the present study evaluates it only in human samples. By retaining observable transcript context, Modulator reveals relationships between transcript architecture and RNA modification that locus-level analyses cannot resolve.

## METHODS

### Samples and sequencing

The HG002 data were the two RNA004 poly(A) libraries released by Oxford Nanopore Technologies. The ZIKV-infected data were three biological replicates each of either mock-infected or ZIKV-infected Huh7 cells. Poly(A) RNA from each replicate was sequenced on one RNA004 flow cell, giving one library per replicate.

### Basecalling and alignment

All libraries were uniformly processed. Raw signal was basecalled with Dorado 2.1.0 using the rna004_sup@v6.0.0 model. The modification basecallers were inosine_m6A_2OmeA, 2OmeG, m5C_2OmeC, and pseU_2OmeU, all version 1, with poly(A) tail estimation. Reads were aligned with minimap2 2.29 in spliced mode with the options -ax splice -uf -k14 -y --secondary=no. The reference was hg38 plus the Epstein-Barr virus genome (NCBI RefSeq NC_007605.1). Alignments were sorted with samtools 1.22.1.

### Per-read modification calls

Modulator runs modkit 0.5.0 pileup on the reads for each fragmentform. By default, a nucleotide on a read is called modified when P(mod) is at least 0.99 and unmodified when P(canonical) is at least 0.8. The valid coverage (Nvalid_cov) is the number of reads with a confident call and the failed count (Nfail) is the number without one. Depth is subsampled to 1,000 reads at deeper sites. A site is kept when two conditions hold: the number of reads that disagree with the reference nucleotide (Ndiff) is at most 3 times the valid coverage (Ndiff ≤ 3 × Nvalid_cov), and the modified count (Nmod) is at least the failed count (Nfail) plus 1 (Nfail + 1 ≤ Nmod).

### Analysis of 5′ read truncation

For **Figure 1a**, we took the 7.7 million reads assigned to fragmentforms that match an annotated hg38 RefSeq transcript exactly. We assumed that the annotated 5′ ends in the hg38 reference transcriptome are accurate. We measured the distance between where reads ended and the annotated 5′ end. We considered a read having reached the annotated 5′ end if it was within 50 nt or less of the annotated 5′ end coordinate.

### Analysis of the impact of read depth on the number of RNA modification sites

For **Figure 1b**, we subsampled the merged HG002 alignments with samtools view -s to 10, 25, 50, 75, and 100% of the 28.2 million mapped reads. We ran modkit pileup with the same settings as previous Modulator runs on each subset and counted the sites with at least 20 reads, greater than 10% modification stoichiometry, Nfail + 1 ≤ Nmod and Ndiff ≤ 3 × Nvalid_cov.

### Identification of full-length read subsets

For **Figure 1d**, we kept only the primary reads whose 5′ alignment end lay within 50 nt of the 5′ end of an overlapping annotated transcript. This kept 7.5 million of 16.6 million reads in library 1 and 8.9 million of 19.9 million in library 2. We ran Modulator on these subset reads with the same settings as the full run.

### Fragmentform assembly and classification

Modulator assigns each read a 3′ end and a junction chain from its alignment. By default, reads with a mapping quality below 10 (meaning the aligner was less than 90% confident of the position) and reads with no splice junction are excluded. Reads with the same junction chain and a 3′ end within 25 nt are grouped, and each group with at least 40 reads becomes a fragmentform. Each fragmentform is classified against the hg38 RefSeq reference transcriptome as an exact match, a new junction chain, a new 3′ end, or a new locus.

### Validation of 3′ ends

For each fragmentform 3′ end, we searched the 40 nt upstream for the 12 poly(A) signals: AAUAAA, AUUAAA, AGUAAA, UAUAAA, CAUAAA, GAUAAA, AAUAUA, AAUACA, AAUAGA, AAAAAG, ACUAAA, and AAGAAA. We also measured the distance to the nearest representative site in the PolyASite 2.0 atlas (release GRCh38.96) on the same strand. As a null, we repeated both tests at a position shifted by a random distance of 1 to 2 kb along the same transcript (shuffled test).

### Reproducibility between HG002 libraries

For each fragmentform, we counted the reads assigned in each HG002 library and scaled each library to reads per million assigned reads. The percent difference is 100 times library 2 minus library 1, divided by the mean of the two. The Spearman correlation was computed on the raw counts.

### Identification of RNA modification stoichiometry differences between fragmentforms

Modulator tests each RNA modification site with at least 20 valid reads in two or more fragmentforms of a gene. It compares the modified and unmodified counts across fragmentforms with the Cochran Mantel Haenszel test, stratified by library (one table per library). When any fragmentform has an expected count below 5 reads in either the modified or the unmodified column, the chi-square approximation is not employed. The p value is instead obtained by simulation: random tables are generated within each library with the same read total per fragmentform and the same modified total as the observed table, the test statistic is computed on each, and the p value is the fraction of random tables whose statistic is at least the observed value. 999 tables are drawn first, extended to 9,999 when fewer than 10 exceed the observed statistic, and to 99,990 when none does. P values are converted to false discovery rates by the Benjamini Hochberg method. A site is considered to have a modification stoichiometry difference when the false discovery rate is at most 0.05 and the largest stoichiometry difference between fragmentforms, weighted by coverage across libraries, is at least 10 percentage points. Each site with differential RNA modification stoichiometry is classified by comparing the most modified and least modified fragmentform.

### Identification of private RNA modification sites

Private sites must have at least 20 reads and a RNA modification stoichiometry of at least 10% in one fragmentform. The site must also lie within a sequence missing from another fragmentform (**Figure 3b**).

### Genotyping in Modulator

Modulator scans the exons of the assembled fragmentforms for candidate SNPs. By default, a candidate requires at least 8 reads in total, at least 4 reads and at least 30% of reads with a nucleotide other than the reference, base quality of at least 20, and mapping quality of at least 10. The reference allele is the hg38 nucleotide at the position and the alternate allele is the most common other nucleotide in the reads. Positions where more than 90% of reads carry the alternate nucleotide are excluded as homozygous, and positions with a third common nucleotide are excluded as multiallelic. By default, candidate SNPs covered by the same molecules are phased into groups of 2 to 4 when at least 4 reads cover every SNP in the group. For each SNP and site carried on the same molecules, Modulator compares the modified fraction between the two alleles with a Cochran Mantel Haenszel test stratified by library, as for the fragmentform test, or with a Fisher exact test when only one library is informative. P values are converted to false discovery rates (FDR) by the Benjamini Hochberg method. A pair is significant when the FDR is at most 0.05 and the modification stoichiometries of the two alleles, weighted by coverage across libraries, differ by at least 10 percentage points. Each pair is placed in a distance class based on the distance between the SNP and the site along the spliced RNA. SNPs that fall on the modified nucleotide itself are excluded from the test and written to a separate table, classified as creating or removing the modifiable nucleotide relative to the reference allele. The reads of all fragmentforms of the gene are pooled for this test. The comparison is stratified by library but not by fragmentform.

### SNP comparison with the HG002 benchmark

SNP candidates were compared with the Genome in a Bottle HG002 GRCh38 v4.2.1 benchmark inside its high-confidence regions on chromosomes 1 to 22 (https://ftp-trace.ncbi.nlm.nih.gov/ReferenceSamples/giab/release/AshkenazimTrio/HG002_NA24385_son/NISTv4.2.1/GRCh38/). A candidate was considered true when a heterozygous benchmark variant had the same position and alternate allele. Phasing was compared with the trio-phased version of the same benchmark.

### Identification of RNA-related changes between conditions

Modulator tests each site within each fragmentform between conditions with a replicate-aware beta-binomial likelihood ratio test, which estimates the variation between replicates and shrinks that estimate across sites. A site within a fragmentform is tested when at least two libraries of each condition have at least 20 valid reads at it. P values are converted to false discovery rates by the Benjamini Hochberg method across all tested sites and modifications. A fragmentform and site pair changes when the false discovery rate is at most 0.05 and the stoichiometry differs by at least 10 percentage points. Sites where the modification itself is read as a SNP by the basecaller are excluded. Modulator similarly identifies changes in fragmentform usage, poly(A) site usage, or junction usage between conditions, with the same test applied to the fraction of a gene’s reads that use each feature in each library, for genes with at least 20 reads per library.

### GLORI

GLORI converts unmodified adenosines to inosine with glyoxal and nitrite while m6A is protected (Liu *et al*. 2023, Shen *et al*. 2024, Sun *et al*. 2025). Methods for GLORI analysis can be found in (Aufgebauer *et al*. 2026). In brief, the same six Huh7 libraries were measured by GLORI, and sites were aligned to hg38. For each site, the m6A stoichiometry is the fraction of unconverted reads. We used a GLORI-reported site when at least two replicates reported it. We define the GLORI change as the ZIKV-infected mean minus the Mock mean.

### Run time

Each dataset was run on one node with the resources as specified in **Supplementary Table 1**. Times are wall clock per stage as reported in the Modulator log outputs.

## DATA AVAILABILITY

Raw direct-RNA sequencing data for HG002 was sourced from the ONT open data repository: https://42basepairs.com/browse/s3/ont-open-data/UHRR_HG002_2026.06. Raw direct-RNA sequencing data for the Huh7 samples is deposited in the European Nucleotide Archive: PRJEB86065. Raw GLORI sequencing data is deposited in the European Nucleotide Archive: PRJEB106901.

## CODE AVAILABILITY

Modulator v2.0.0 is available on GitHub (https://github.com/Theo-Nelson/modulator) and archived at Zenodo (https://doi.org/10.5281/zenodo.23381616).

## ACKNOWLEDGEMENTS

C.E.M. was supported by the Katherine and John Bleckman Foundation, and the NIH (U54AG089334, R01AI125416). Research on m6A in the Horner lab has been supported by Burroughs Wellcome Fund and National Institutes of Health (NIH) grant R01AI125416. T.M.N. was supported by a Medical Scientist Training Program grant from the National Institute of General Medical Sciences of the National Institutes of Health under award number: T32GM152349 to the Weill Cornell/Rockefeller/Sloan Kettering Tri-Institutional MD-PhD Program. T.M.N. was also supported with computational resources from the National Science Foundation ACCESS Allocation Request BIO240371. T.M.N. gratefully acknowledges use of the research computing resources of the Empire AI Consortium, Inc, with support from Empire State Development of the State of New York, the Simons Foundation, and the Secunda Family Foundation (10.1145/3708035.3736070). T.M.N. also acknowledges the Weill Cornell Medicine High-Performance Computing Core for providing computational resources and support that contributed to the results reported in this paper. RRID:SCR_028969. Additionally, this material is based upon work supported by the National Science Foundation Graduate Research Fellowship (C.J.A), and the CMB training grant T32GM142605 (C.J.A). E.M.V. was supported by the Weill Cornell ACCESS Summer Internship Program.

## AUTHOR CONTRIBUTIONS

Conceptualization, T.M.N., C.J.A., S.M.H., and C.E.M.; Methodology, T.M.N.; Software, T.M.N. and M.G.; Validation, T.M.N. and C.J.A.; Formal analysis, T.M.N., M.G., N.H., E.M.V., and E.A.S.; Investigation, T.M.N., C.J.A., P.C., J.P., and K.R.; Resources, T.M.N., C.J.A., P.C., J.P., K.R., S.M.H., and C.E.M.; Data curation, T.M.N.; Writing – original draft, T.M.N. and M.G.; Writing – review & editing, T.M.N., M.G., S.M.H., and C.E.M.; Visualization, T.M.N. and M.G.; Supervision, S.M.H. and C.E.M.; Project administration, S.M.H. and C.E.M.; Funding acquisition, S.M.H. and C.E.M.

## TERMS

### Direct RNA sequencing

A nanopore sequencing method that reads native RNA molecules one at a time, from the 3′ end toward the 5′ end.

### Read

The sequence and modification calls reported from one RNA molecule by the Nanopore basecaller.

### 5′-truncated read

A read that terminates before the 5′ end of its transcript.

### Library

The reads from one sequencing run of one sample on one flow cell. HG002 was sequenced on two flow cells, ‘library 1’ and ‘library 2’.

### Reference transcriptome

The set of annotated transcripts for a genome. This work uses the hg38 RefSeq annotation.

### Splice junction

The point where two exons are joined after the intron between them is removed.

### Donor and acceptor

The two boundaries of an intron. The donor is the 5′ boundary and the acceptor is the 3′ boundary.

### Junction chain

The ordered list of splice junctions in a read or a fragmentform.

### Fragmentform

The 3′ end of a transcript together with the junction chain upstream of it. Every read is assigned to a single fragmentform, whether or not it reaches the 5′ end.

### Poly(A) site

The position where a transcript is cleaved and the poly(A) tail is added. It is located at the 3′ end of the transcript.

### Poly(A) signal

A six-nucleotide sequence, such as AAUAAA, within 40 nt upstream of a poly(A) site, which directs the cleavage.

### Alternative poly(A) sites

Two poly(A) sites in the same last exon. The proximal site is closer to the 5′ end and the distal site is farther from it.

### Intronic poly(A) site

A poly(A) site inside an intron, so that the transcript ends before the next exon.

### RNA modification

A chemical change to a nucleotide in RNA. This work includes all modifications currently called by the Nanopore basecaller, including N6-methyladenosine (m6A), inosine, pseudouridine (pseU), 5-methylcytidine (m5C), and the four 2′-O-methyl modifications (2′OmeA, 2′OmeC, 2′OmeU, 2′OmeG).

### Stoichiometry

The fraction of molecules that carry the modification at one position.

### Position

A single nucleotide in the reference genome.

### Site

A position at which a modification is measured.

### Private site

A site present in one fragmentform and absent from another, because the other fragmentform splices it out or ends before it.

### Shared site

A site present in both fragmentforms that are compared.

### Difference and change

A difference in RNA modification stoichiometry is between fragmentforms or between alleles within one sample. A change in RNA modification stoichiometry is between conditions.

### Unexplainable

A difference between fragmentforms that only the 5′ end information could explain.

### SNP

A single nucleotide polymorphism, a position where the two copies of a gene carry different nucleotides.

### Allele

One of the two versions of a gene, or of a SNP, in a diploid cell. The reference allele matches the reference genome and the alternate allele does not.

### Phasing

Determining which SNP alleles lie on the same molecule.

### DRACH motif

The five-nucleotide sequence preferred by the enzyme that adds m6A (D = A, G, U; R = A, G; H = A, C, U).

### Exon junction complex

A group of proteins deposited upstream of each splice junction during splicing.

### Condition

A treatment state. In this work, mock infection (’Mock’) or Zika virus infection (’ZIKV-infected’).

### GLORI

A chemical method that measures m6A stoichiometry at single nucleotides, independent of nanopore sequencing.

## SUPPLEMENTARY INFORMATION

**SUPPLEMENTARY TABLE 1.** Run time and memory per dataset.

| Dataset | Libraries | Reads in BAM | Mapped reads | Reads assigned to fragmentforms | Fragment forms | Node | Peak memory | Total wall time |
| --- | --- | --- | --- | --- | --- | --- | --- | --- |
| HG002 | 2 | 36.3 M | 28.2 M | 16.1 M | 23,746 | 48 cores, 300 GB | 24.9 GB, assembly | 8.3 h |
| Huh7 Mock and ZIKV-infected | 3 + 3 | 135.7 M | 103.9 M | 69.4 M | 59,699 | 48 cores, 300 GB | 58.9 GB, assembly | 18.6 h |

Stage times for HG002 were as follows. Fragmentform assembly took 27 min, read statistics 6 min, and modification calls 4.2 h. Site aggregation took 12 min, fragmentform tests 1 min, classification 5 min, and the multi-gene filter 12 min. Genotyping took 2.5 h, poly(A) tails 26 min, and the report 2 min. Times are wall clock with 6 parallel jobs.

**SUPPLEMENTARY TABLE 2.**
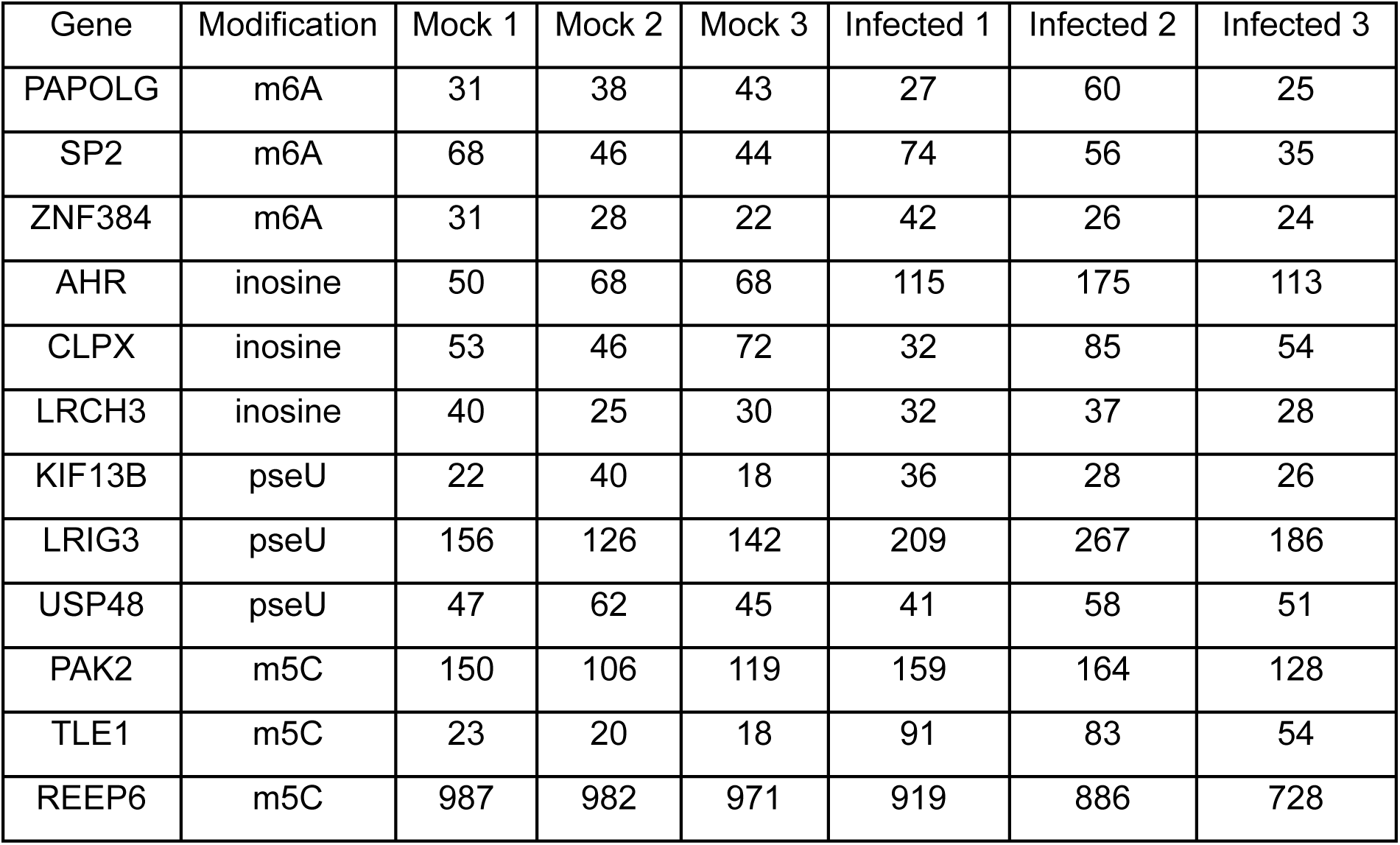
Reads per library at the 12 examples of Figure 7c.

